# Dynamic population coding of social novelty in the insular cortex

**DOI:** 10.1101/2024.04.01.587524

**Authors:** Masaaki Sato, Eric T. N. Overton, Shuhei Fujima, Toru Takumi

**Affiliations:** Department of Neuropharmacology, Hokkaido University Graduate School of Medicine, Kita, Sapporo, Japan; RIKEN Center for Brain Science, Wako, Japan; Department of Physiology and Cell Biology, Kobe University School of Medicine, Chuo, Kobe, Japan; RIKEN Center for Biosystems Dynamics Research, Chuo, Kobe, Japan

**Author notes:** For correspondence (TT).

## Abstract

The familiarity of socially interacting peers has a profound impact on behavior^1–3^, but little is known about the neuronal representations distinguishing familiar from novel conspecifics. The insular cortex (IC) regulates social behavior^4–9^, and our previous study revealed that neurons in the agranular IC (aIC) encode ongoing social interactions^10^. To elucidate how these neurons discriminate between interactions with familiar and novel conspecifics, we monitored neuronal activity in mice by microendoscopic calcium imaging during social recognition memory (SRM) and linear chamber social discrimination (LCSD) tasks. In the SRM task, repeated interactions with the same target activated largely nonoverlapping cells during each session. The fraction of cells associated with social investigation (social cells) decreased as the subject repeatedly interacted with the same target, whereas substitution of a second novel target and subsequent exchange with the first familiar target recruited more new social cells. In the LCSD task, the addition of a novel target to an area containing a familiar target transiently increased the number of cells responding to both targets, followed by an eventual increase in the number of cells responding to the novel target. These results support the view that the aIC dynamically encodes social novelty, rather than consistently encode social identity, by rapidly reorganizing the neural representations of conspecific information.

## Main text

In animals, recognition of whether interacting conspecifics are familiar or novel profoundly influences social behavior^1–3^. This recognition is thought to be supported by memory and novelty detection, but the underlying neural representations remain poorly understood. We formulated two contrasting hypotheses to explain how agranular insular cortex (aIC) neurons discriminate familiar from novel conspecifics during social interactions. A “consistent coding” model posits that unique subsets of cells encode interactions with individual conspecifics (**Figure 1A, left**). Alternatively, a “dynamic coding” model predicts that repeated interactions with the same individual activate largely nonoverlapping cells each time (**Figure 1A, right**). The IC is involved in both memory^9,11,12^ and salience detection^13,14^, so both scenarios are possible^4^. To distinguish between these alternative models, we monitored the activities of aIC pyramidal neurons in mice expressing the genetically-encoded calcium indicator GCaMP6f during a social recognition memory (SRM) task^10^ (**Figure 1B and C**). A male subject mouse with a microendoscope attached to its head was allowed to explore a linear chamber containing a wire cage with either no mouse (Emp), stimulus mouse 1 (S1), or stimulus mouse 2 (S2). In the first phase (familiarization), the subject mouse was allowed to interact with S1 three times at 0 h (S1-0), 1 h (S1-1), and 24 h (S1-24) to examine the short- and long-term representational stability of cells responsive to social interactions (social cells, SCs). In the second phase (discrimination), the subject consecutively interacted with the now familiar S1 (S1-24, also part of the familiarization phase), the unfamiliar S2 (S2-24), and S1 again (S1-24-R) to examine representational changes between novel and familiar individuals.

**Figure 1.**
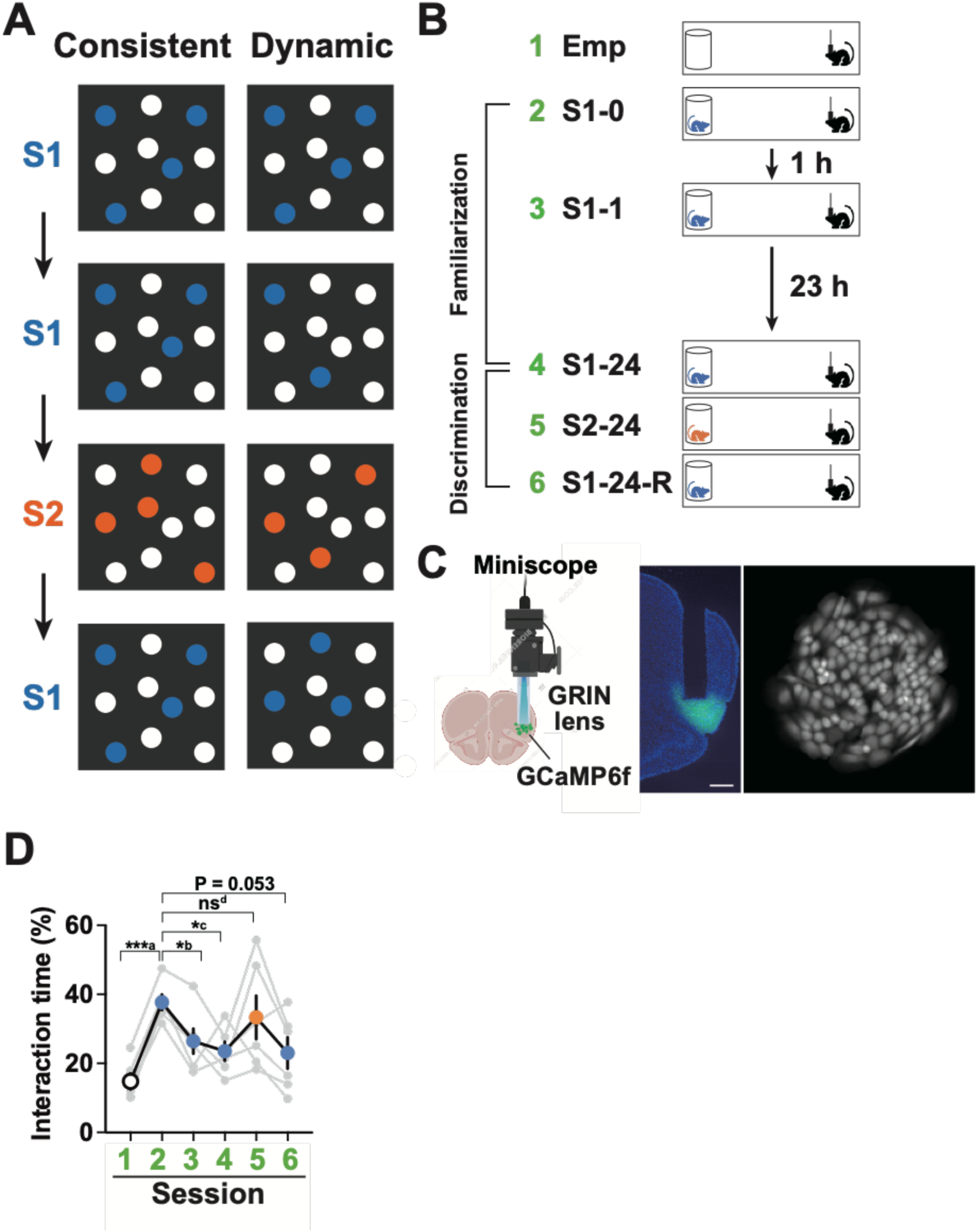
**Proposed models and the SRM task. A**. Two models for encoding social interactions with familiar and novel conspecifics by aIC neurons. In the consistent model, repeated interactions with the same stimulus mouse (S1) activate the same subset of neurons, whereas an interaction with a different stimulus mouse (S2) activates a different subset of neurons (left). In the dynamic model, the subset of cells activated by repeated interactions with S1 is not fixed (right). **B**. SRM task. The task consists of a familiarization phase and a discrimination phase. In each session, a subject mouse was allowed to explore a test chamber containing a wire cage for 5 min. In Session 1 (Emp), the wire cage was empty. In Session 2 (S1-0) conducted shortly after Session 1, the cage contained the first stimulus mouse (S1). This protocol was repeated 1 h (Session 3, S1-1) and 24 h (Session 4, S1-24) after Session 1. Session 4, which is also the first session of the discrimination phase, was immediately followed by an interaction session with the novel second stimulus mouse (Session 5, S2-24) and a session of re-exposure to S1 (Sessions 6, S1-24-Re). **C**. Microendoscopic calcium imaging. Left: GCaMP6f-labeled aIC neurons were imaged using a miniaturized head-mounted fluorescence microscope (Miniscope) through a chronically implanted GRIN lens. Middle: A fluorescence image of a coronal section showing GCaMP6f-labeled aIC neurons and the outline of the GRIN lens (scale bar = 500 µm). Right: A reconstructed FOV showing 147 aIC neurons identified by the CNMFe algorithm. **D**. Percentage time spent by the subject mouse investigating the interaction target in the wire cage during each session. ***a, *P* < 0.0001; *b, *P* = 0.013, and *c, *P* = 0.013; ns^d^, *P* = 0.445 by repeated-measures one-way ANOVA with post-hoc Holm–Sidak tests. n = 6 mice.

Repeated exposure to S1 significantly reduced the interaction time during S1-1 and S1-24, while the interaction time with S2 during S2-24 reversed this reduction and the interaction time with S1 during S1-24-R returned to the same level as S1-24 (**Figure 1D**). Changes in the fluorescence of GCaMP6f-labeled neurons (total 456 cells from 6 mice, 50–96 cells/mouse) revealed that the fraction of neurons activated by social interactions (termed Social-ON cells, **Figure 2A**) was highest at S1-0 and decreased thereafter during S1-1 and S1-24 (**Figure 2B**). During S2-24, the activated fraction did not decrease further but declined again during S1–24-R. The fraction of neurons inhibited by social interactions (termed Social-OFF cells) was low and did not change significantly during repeated S1 exposure (**Figure 2C**). Thus, we focused on Social-ON cells and hereafter refer to them simply as SCs.

**Figure 2.**
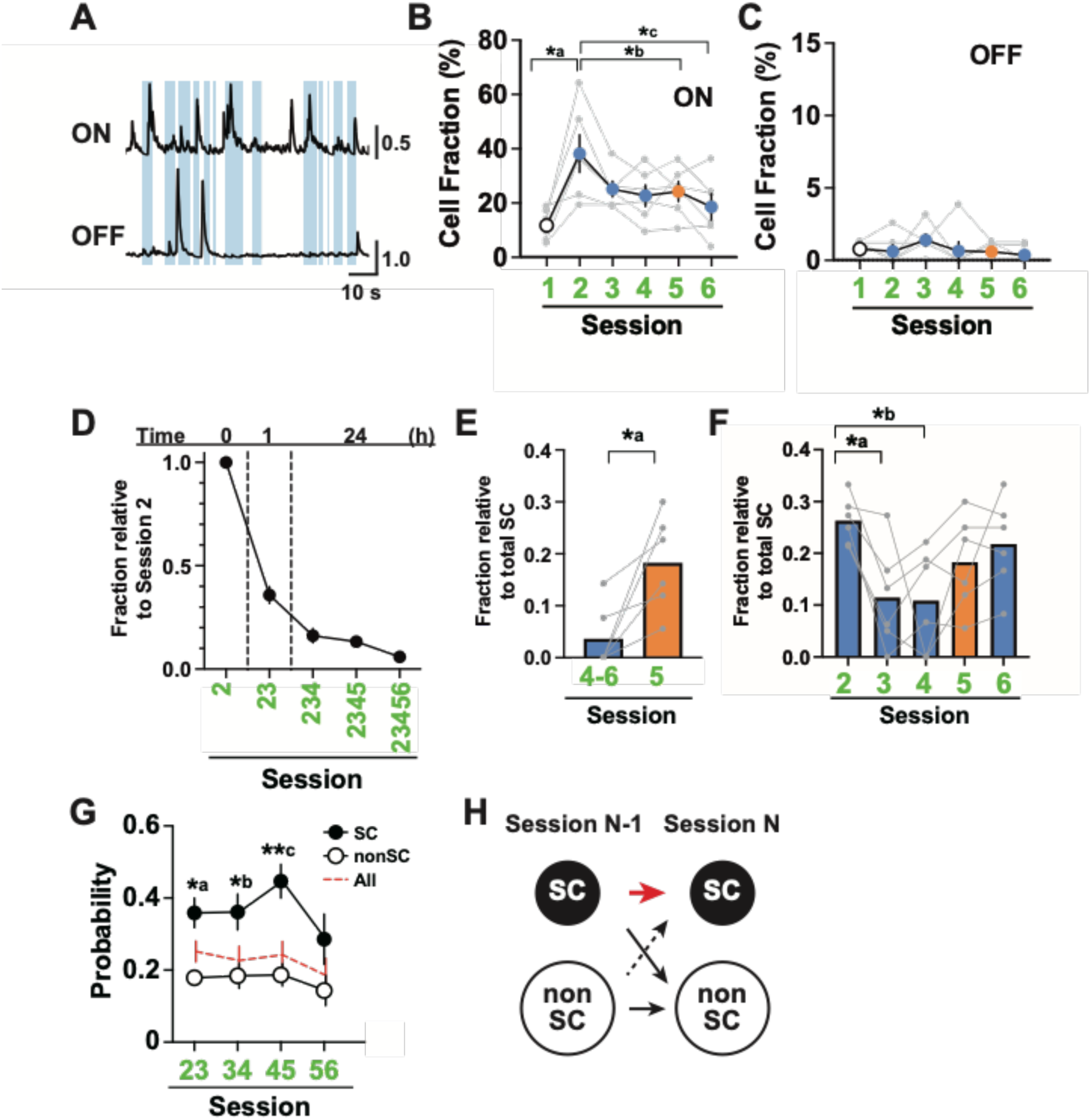
Calcium imaging during the SRM task. A. GCaMP fluorescence traces from a Social-ON cell and a Social-OFF cell. Social interaction periods are indicated in blue. **B**. The fractions of Social-ON cells relative to the total number of identified neurons during each session. n = 6 mice. *a, *P* = 0.038; *b, *P* = 0.041; *c, *P* = 0.033 by repeated-measures one-way ANOVA with post-hoc Holm–Sidak tests. **C**. Fractions of Social-OFF cells relative to the total number of identified neurons during each session. n = 6 mice. **D**. Fractions of cells that acted as SCs during subsequent consecutive sessions relative to the number acting as SCs during Session 2. n = 6 mice. **E**. Fractions of cells acting as SCs only in Sessions 4 and 6 (4-6) and only in Session 5 (5) relative to the total number of SCs identified in the relevant session. *a, *P* = 0.031 by Wilcoxon matched-pairs signed rank test. n = 6 mice. **F**. Fractions of cells acting as SCs only during the indicated session relative to the total number of SCs identified in the relevant session. *a, *P* = 0.025; *b, *P* =0.032 by repeated-measures one-way ANOVA with post-hoc Holm–Sidak tests. n = 6 mice. **G.** Probabilities of SCs (filled) and nonSCs (open) during Session N-1 that also acted as SCs during the next Session N. The red dashed line delineates the data for all cells. The two digits on the X-axis represent session pairs considered. n = 6 mice. *a, *P* = 0.016, *b, *P* = 0.034; **C, *P* = 0.005, SC vs. nonSC by repeated-measures two-way ANOVA with post-hoc Holm–Sidak tests. **H**. A model of SC stability over consecutive sessions. SCs in the previous session were more likely than nonSCs to be SCs in the current session. The red arrow and dashed arrow indicate augmentation and weakening of the indicated processes, respectively.

We examined the representational stability of SCs (see Methods). In the familiarization phase, the fraction of SCs stable for 1 h and 24 h was 35.9 ± 4.1 % and 16.2 ± 3.4 %, respectively (**Figure 2D**). Fifty-eight of 110 cells active during S1-1 (52.7%) were also active during S1-0, whereas the remaining 52 cells (47.3%) active during S1-1 exhibited newly acquired SC activity. Furthermore, very few SCs (only 5.9% ± 2.4%) were active during all five social interactions (from S1-0 through S1-24-R, Figures 2D and Figure supplement 1).

Next, we analyzed whether distinct subsets of SCs are activated during interactions with different social targets. After 24 h of familiarization, the fraction of neurons that responded as SCs only during interaction sessions with S1 (Sessions 4 and 6) was significantly lower than the fraction that responded as SCs during the interaction session with S2 (Session 5) (**Figures 2E and Figure supplement 1**). We then determined the fractions of cells that were active during only one of the five interaction sessions. The fractions active only during S1-1 or S1-24 were significantly smaller than the fraction active during S1-0 (**Figure 2F**). In contrast, the fraction of cells responding as SCs only during S2-24 or S1-24-R did not differ significantly from the fraction active only during S1-0. We further examined whether SCs in one session tended to be SCs in subsequent sessions. The probability of an SC during Session N-1 also acting as an SC during Session N was higher than that of a nonSC during Session N-1 acting as an SC during Session N (**Figure 2G**). Moreover, the probability of SC recurrence was higher for interactions with a novel target (transition between Session 4 and 5), suggesting that the acquisition of SC activity is not randomly assigned. Therefore, although representations drifted, SCs during one session were more likely than nonSCs to act as SCs again during a subsequent session (**Figure 2H**).

Results thus far suggest that the fraction of aIC neurons acting as SCs may depend on the time spent interacting with social targets (**Figures 1B and 2B**). The fraction of SCs and the total duration of social interaction were moderately to highly correlated for each individual mouse (r = 0.33–0.87, n = 6 mice; **Figure supplement 2A**). However, the correlations markedly weakened when calculated using the data pooled from all six mice (**Figure supplement 2B–C**). Further, the positive correlations observed in the subject-wise analysis were in striking contrast to the widely varying correlations (slopes of linear regression lines) obtained from session-wise analysis (**Figure supplement 3**). This implies that SC population size predicts the total length of social interaction at the within-subject level but not at the across-subject level.

Next, we imaged aIC neuron activity while the mice performed an LCSD task (**Figure 3A**). Wire cages placed at each end of a linear chamber contained either no mouse (Emp), an inanimate object (Obj), or a stimulus mouse (S1, S2, or S3) as indicated. Following the baseline session with empty cages (Emp1 and Emp2), the subject was allowed to investigate stimulus pairs Obj:S1, S2:S1, and S1:S3 successively according to the arrangement shown in Figure 3A. Times spent exploring the two empty cages were comparable during the baseline session (**Figure 3B**) while the time spent investigating S1 was significantly longer than that investigating Obj during the first social session (**Figure 3C**). While the subject mouse spent only slightly more time investigating S2 than S1 during the second session (**Figure 3D**), it spent significantly more time investigating S3 than S1 during the last session (**Figure 3E**), suggesting that familiarity with S1 gradually developed over the last two sessions.

**Figure 3.**
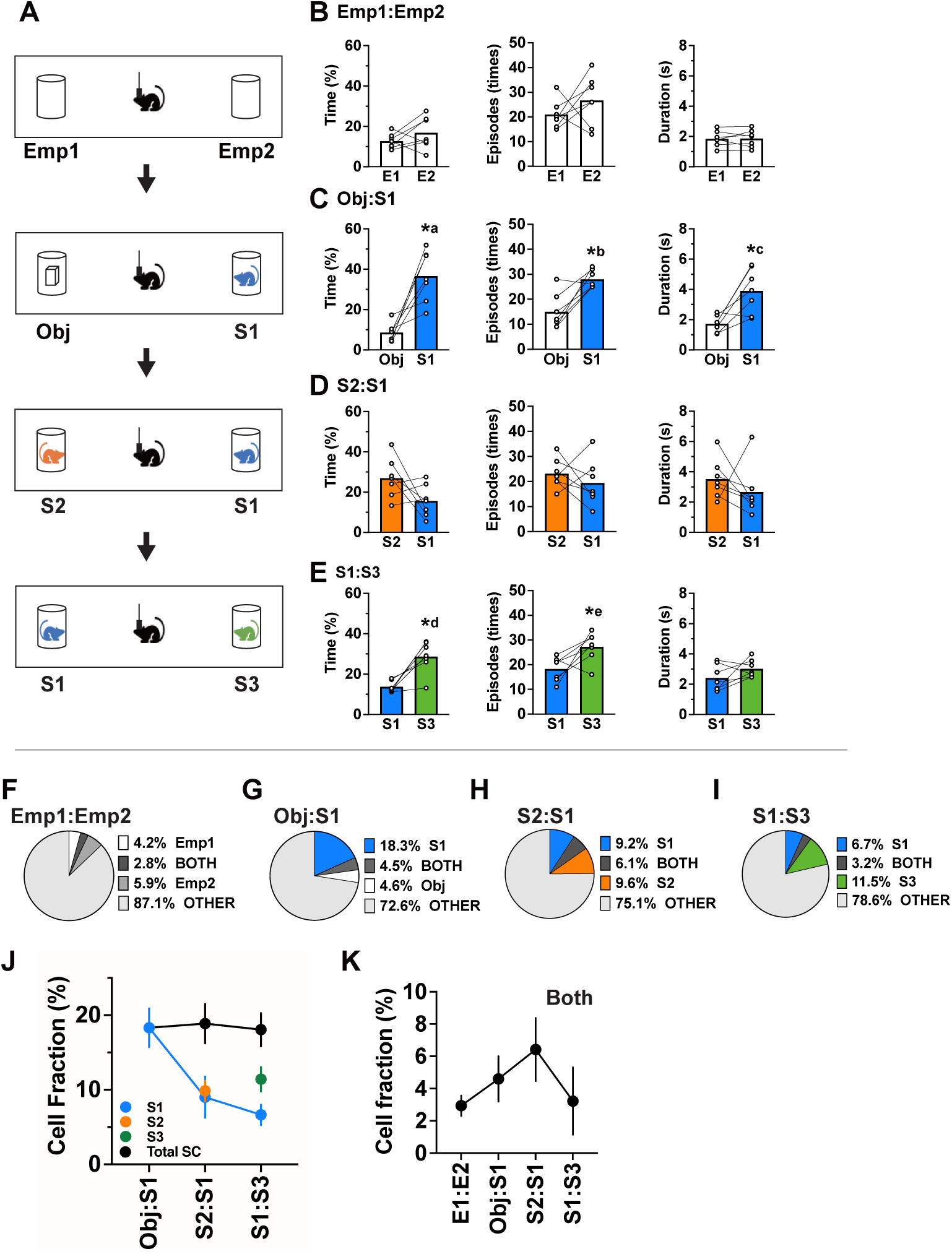
Behavioral analysis and calcium imaging during the LCSD task. A. LCSD task. A subject mouse with a microendoscope attached to its head interacted with various targets contained within two wire cages at each end of a linear chamber. The task consisted of four sessions. The two wire cages contained no targets (Emp1:Emp2), an inanimate object and stimulus mouse 1 (Obj:S1), stimulus mouse 2 and S1 (S2:S1), and S1 and stimulus mouse 3 (S1:S3) in that order. The position of S1 was maintained at the same end of the chamber in the second and third sessions, whereas it was moved to the other end in the fourth session. The sessions were 5 min long and conducted consecutively without intervals. **B–E** The fraction of time spent investigating each target (left), the number of social interaction episodes (middle), and average duration of interaction episodes (right) in Emp1:Emp2 (B), Obj:S1 (C), S2 :S1 (D), and S1:S3 (E) sessions. *a, *P* = 0.016; *b, *P* = 0.031; *c, *P* = 0.031; *d, *P* = 0.016; *e, *P* = 0.047 by Wilcoxon matched-pairs signed rank test. n = 7 mice. **F–I** The fractions of cells encoding the interaction with each target during Emp1:Emp2 (F), Obj:S1 (G), S2:S1 (H), and S1:S3 (I) sessions. Other includes small numbers of Social-OFF cells. **J.** Changes in the fractions of S1-responsive SCs (blue) and total SCs (black) across sessions. Total SCs in S2:S1 and S1:S3 sessions represent the sum of SC fractions against S1 and S2 (orange) and those against S1 and S3 (green), respectively. n = 7 mice. **K**. Change in the fraction of SCs that responded to both interaction targets across sessions. n = 7 mice.

We then calculated the fraction of SCs that responded to interactions with each target during all session (969 cells from 7 mice, 100–170 cells/mouse, **Figure 3F–I and Figure supplement 4**). For the three repeated exposures to S1, the fraction of S1-responsive cells was highest (18.3%, 177 cells) during the first session and decreased monotonically thereafter (**Figure 3F–J**). Furthermore, the fraction of cells that responded to novel mouse S2 or S3 was no less than that responding to the previously encountered S1 (**Figure 3H and I**). The overall fraction of responding cells did not double even when the number of social targets was doubled by the addition of S2 or S3 (**Figure 3J**), suggesting limited allocation of aIC neurons to social representations. Notably, the fraction of cells that responded to both targets transiently increased during the S2:S1 sessions (6.1%, 59 cells) and returned to the previous level during the S1:S3 sessions (**Figure 3K**). This increased fraction of cells with broad response selectivity paralleled the poor discrimination between S2 and S1 at the behavior level as evidenced by similar investigation times (**Figure 3D**).

Finally, we analyzed the transitions of functional cell categories between consecutive sessions. An analysis of 177 cells that responded to S1 during Session 2 revealed that more cells shifted response to novel S2 during Session 3 than continued responding to familiar S1 (**Figure 4A**). Further, the fraction of S2-responsive cells during Session 3 that also responded to novel S3 during Session 4 was larger than the fraction responding to familiar S1 during Session 4 (**Figure 4B, left**). Similarly, the fraction of S1-responsive cells during Session 3 that responded to novel S3 during Session 4 was larger than the fraction that continued responding to familiar S1 during Session 4 (**Figure 4B, middle**). Cells that responded to both S2 and S1 during Session 3 also exhibited a similar tendency to shift response toward novel S3 during Session 4 (**Figures 4B, right and Figure supplement 4**). Moreover, only 0.4% of cells (4 of 969) persistently responded to S1. Together, these results demonstrate that SCs tend to represent interactions with novel targets rather than familiar targets during subsequent sessions (**Figure 4C**).

**Figure 4.**
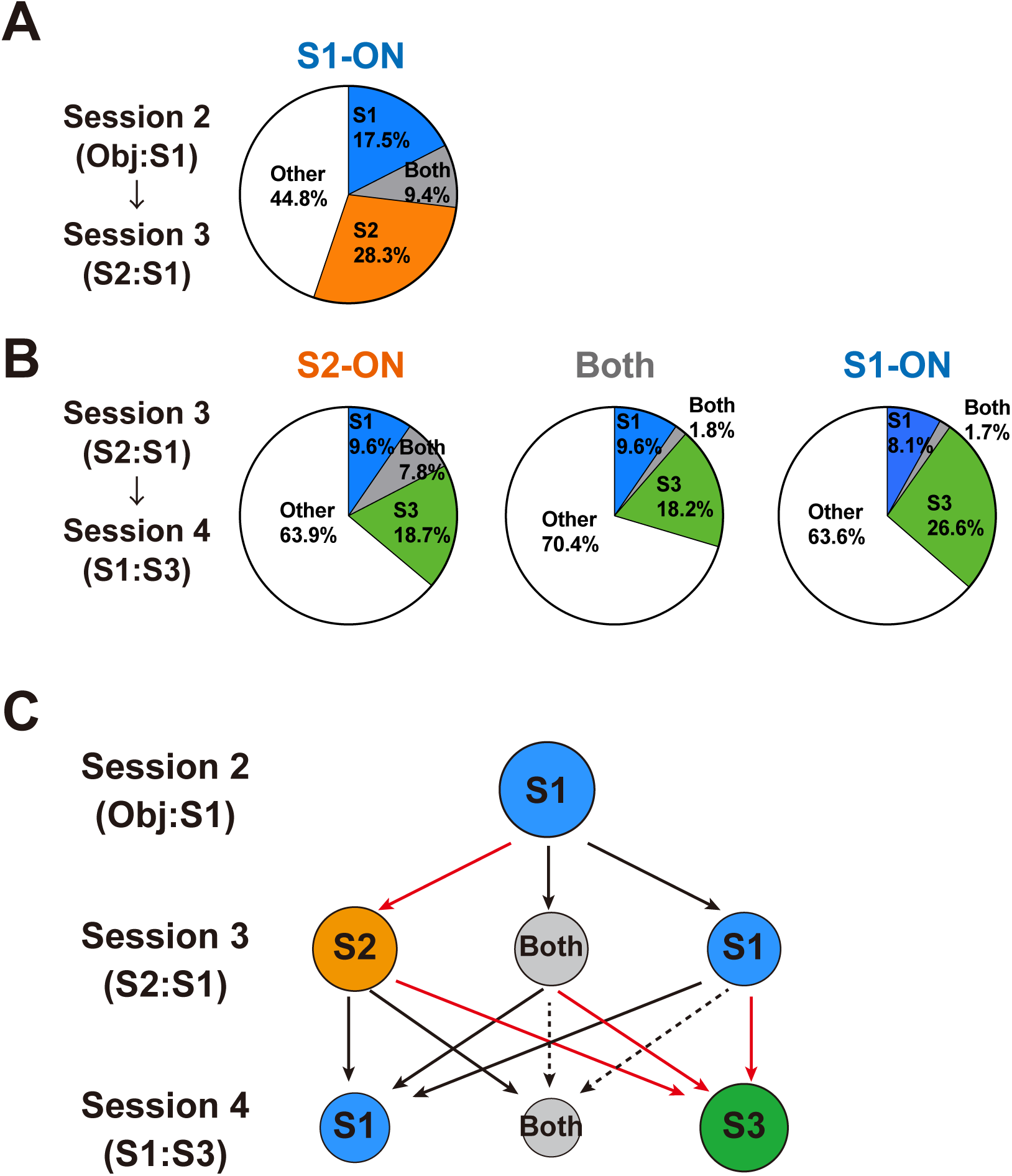
Transition of functional cell categories across sessions in the LCSD task. A. Pie chart showing the average fractions of S1-responsive SCs during Session 2 that also acted as SCs in response to the indicated target. n = 7 mice. **B**. Pie charts showing average fractions of S2-responsive SCs (left), S1-responsive SCs (right), and SCs that responded to both S1 and S2 (middle) during Session 3 and also responded to the indicated target in Session 4. n = 7 mice. **C.** Transitions in the representations of social interaction targets across sessions during the LCSD test. Red and dotted arrows represent enhancement and reduction of the indicated process, respectively.

Results of the SRM and LCSD tasks support the idea that SCs in aIC encode social novelty rather than social target identity through dynamic changes in target responsivity. Such unique social novelty coding in aIC may allow this structure to effectively broadcast salience signals to downstream networks that enable flexible social behavior^2,5–9,15^. In rodents, information about conspecifics such as social identity and familiarity^16^ as well as relevant spatial and sexual information^17,18^ is encoded by distinct brain regions. Salient auditory stimuli have been reported to activate the anterior IC and subsequently suppress default mode network nodes^14^. Since novelty is an important element of saliency, our findings imply that the rodent aIC detects salient stimuli in both the nonsocial and social domains. Important questions such as how social interaction leads to the development of “social fields” in a subset of neurons remain to be addressed in future investigations. Given findings from other brain regions, over-representation of a novel conspecific in the aIC could be mediated by processes involving dedicated synaptic plasticity^19,20^ and neuromodulation^21,22^.

### Materials and methods Ethics

All experiments were performed according to guidelines and protocols approved by the RIKEN Animal Experiments Committee. The protocol complied with the Fundamental Guidelines for the Proper Conduct of Animal Experiment and Related Activities in Academic Research Institutions under the jurisdiction of the Ministry of Education, Culture, Sports, Science and Technology of Japan.

### Mice

Adult male C57BL/6J mice were purchased from Japan SLC (Hamamatsu) or obtained from in-house breeding. All mice were maintained under a 12 h: 12 h light: dark cycle (lights on at 8:00 a.m.) with ad libitum access to food and water. Surgeries were performed at 8 weeks of age or older. Mice that received surgery were maintained under a reverse light: dark cycle (lights off at 8:00 a.m.) and allowed to recover for at least 2 weeks before experimentation. Behavioral experiments were performed at 10 weeks of age or older during the dark phase of the circadian cycle.

### Surgery

Surgery was performed essentially as described previously^10^. Briefly, mice were microinjected with an adeno-associated viral (AAV) vector encoding the calcium indicator GCaMP6f under control of a neuron-specific calcium/calmodulin-dependent protein kinase II (CaMKII) promoter and implanted with a gradient refractive index (GRIN) lens in the unilateral aIC on the same day. The baseplate for a miniaturized head-mounted fluorescence microscope (miniscope, nVista, Inscopix) was then affixed onto the skull on a later day. In detail, mice were first anesthetized with isoflurane (2% in oxygen; flow rate, 0.5 l/min) and placed in a stereotactic frame (Kopf Instruments). After removing a piece of scalp, a small craniotomy was performed above the right aIC. A 35- gauge needle attached to a microsyringe (World Precision Instruments) was directed toward the right aIC (1.94 mm anterior, 2.2 mm lateral, and 3.45 mm deep relative to bregma), and 500 nl of virus solution (AAV5-CaMKII-GCaMP6f-WPRE-SV40, 1.0 × 10^14^ GC/mL, Penn Vector Core) was microinjected over 5 min. A GRIN lens (0.5 mm diameter, 4.0 mm length, GLP-0540, Inscopix) attached to a lens implant kit (ProView, Inscopix, Palo Alto, CA) was inserted slowly 200 µm dorsal to the AAV injection site and affixed to the skull with dental cement. Four weeks later, the baseplate (Inscopix) attached to a miniscope was positioned above the implanted lens on the skull under isoflurane anesthesia using an adjustable gripper (Inscopix). The microscope and baseplate were lowered toward the top of the lens until GCaMP fluorescence signals were observed. The baseplate was then affixed to the skull using dental cement. Mice were housed individually after AAV vector microinjection and GRIN lens implantation until the end of the experiment.

### Behavior

To investigate changes in neural representations associated with familiarity and discrimination of conspecifics, SRM and LCSD tasks were conducted. Both tasks were conducted in an acrylic linear chamber (60 cm long, 10 cm wide, and 21 cm high) divided into a central chamber (40 cm long) and two flaking test chambers (10 cm long each) as needed using plastic dividers. Subject mice were handled for at least 5 days before the behavioral experiments. Mice were also connected to the miniscope and allowed to habituate to the experimental environment for 20 min per day on two consecutive days before testing.

For the SRM task, one test chamber was walled off with a divider. A wire cage (an inverted wire pencil cup) was placed on the opposite side of the chamber. The task consisted of six sessions over two days that tested short-term (1 h) and long-term (24 h) social recognition memory (**Figure 1B**). In the first session, a subject mouse was placed in the chamber and allowed to move freely for 5 min with no stimulus mouse in the wire cage (Emp session). In the second session, the first male stranger stimulus mouse (S1) was placed in the wire cage, and the subject mouse was allowed to freely interact with S1 for an additional 5 min (S1-0 session). During the third session conducted 1 h later, this interaction with S1 was repeated (S1-1 session) using the same chamber arrangement as in the second session. Twenty-four hours after the first session, the fourth, fifth, and sixth sessions were conducted as follows. The subject mouse was allowed to interact with S1 in the wire cage for 5 min (S1-24 session), followed by interaction with the second novel male stimulus mouse (S2, S2-24 session) and finally re-exposure to S1 (S1-24-R session). The LCSD task was designed to examine the familiarity and discriminability of multiple novel conspecifics and was performed in a linear chamber with two wire cages placed on both ends. This task consisted of four consecutive 5-min sessions conducted over one day (**Figure 3A**). In the first session, a subject mouse was allowed to explore the linear chamber with two empty wire cages placed at each end (Emp1:Emp2 session). In the second session, an inanimate object and the first male stranger stimulus mouse (S1) were placed in separate wire cages, and the subject mouse was allowed to freely explore the environment (Obj:S1 session). In the third session, the inanimate object was replaced with the second male stranger stimulus mouse (S2), while S1 remained in the same wire cage (S2:S1 session). Again, the subject mouse was allowed to freely explore the chamber for 5 min. In the last session, S1 was moved to the other wire cage, the third male stranger stimulus mouse (S3) was introduced into the cage previously containing S1, and the subject mouse was allowed to freely explore this environment (S1:S3 session). The location of S1 was moved between the third and last sessions to control for the effect of location. The location of S1 in the second and subsequent sessions was counterbalanced across subjects.

Behavior was recorded throughout the test using a video camera (logicool). Behavioral recordings were synchronized with calcium imaging by simultaneously recording the frame counter of the imaging movie within the behavior video screen. Different stranger stimulus mice were used in the SRM and LCSD tasks. In both tasks, the wire cages were wiped with 70% ethanol after every session, and the entire apparatus was wiped before sessions with a new subject mouse.

The experimental schedule consisted of two days of SRM sessions followed by one day of LCSD sessions. A total of 7 subject mice were examined in SRM and LCSD experiments. Data from all 7 mice were included in the analysis of LCSD experiments. The SRM results for one mouse were unintentionally lost, so data from only 6 mice were included in the analysis.

### Microendoscopic calcium imaging in freely moving mice

The activities of GCaMP6f-labeled aIC neurons were imaged during social behavior tasks using a miniscope and the implanted GRIN lens^10^ (**Figure 1C**). Before the experiment, mice were briefly anesthetized with 2% isoflurane to remove the baseplate cover from the head. The miniscope was then attached to the baseplate. Mice were allowed to recover from anesthesia for at least 20 min before the behavioral task.

Movies of time-varying changes in GCaMP6f fluorescence signals were acquired using nVista (Inscopix) at a resolution of 1440 × 1080 pixels and a rate of 20 frames/s. The LED power of the microscope was set between 0.7 and 1.2 mW. Histological examinations conducted after the experiments verified that the implanted end of the GRIN lens was appropriately positioned over the aIC in all seven subject mice.

### Histology

Mice were administered an overdose of isoflurane and transcardially perfused with 0.2 M phosphate-buffered saline (PBS) followed by 4% paraformaldehyde (PFA) in PBS. Dissected brains were post-fixed in 4% PFA and sliced into 50 µm-thick sections using a vibratome (Leica VT1200 S). Sections were mounted on glass slides and coverslipped with mounting medium containing DAPI (Vectashield, H1200). Fluorescence images were obtained using an epifluorescence microscope (Keyence BZ 9000).

### Data analysis

The behavior of each subject mice was manually classified by visual inspection of the videos after experiments. In both the SRM and LCSD experiments, we defined the interaction period as the time actively exploring the target across the wire cage by nasal contact and sniffing. The remaining periods were classified as nonsocial.

Calcium imaging movies were processed using Inscopix Data Processing Software (IDPS) as described previously^10^. In brief, videos were spatially downsampled by a factor of 2, motion corrected, and normalized as described previously^10^, yielding fluorescence changes (ΔF/F0) for each cell in the field of view (FOV). Individual cells were then identified using an extended constrained nonnegative matrix factorization algorithm for microendoscopic data (CNMFe; parameters, gSig = 3, gSiz = 20, min_pnr = 15, min_corr = 0.8), followed by human visual comparison against the ΔF/F0 video. For analysis of the SRM experiments, movies recorded consecutively were concatenated (i.e., Emp and S1-0 movies were concatenated into one movie and S1-24, S2-24 and S1-24-R movies were concatenated into another) for cell identification. After cell identification, cells that appeared across movies at different time points were verified by longitudinal cell registration (LCR) in IDPS (parameter, minimum correlation = 0.7). In the LCSD experiments, cells were identified from single concatenated movies that spanned all four sessions, so LCR was not necessary. Overall, 121 ± 28 cells/FOV were identified after visual verification (mean ± SD, n = 19 concatenated movies from 7 mice). Calcium transients were detected using an event detection algorithm in IDPS (parameters, event smallest decay time = 0.20 s, event threshold factor = 5).

Neurons exhibiting activity correlated or anticorrelated with social behavior were identified by statistical testing of cosine similarity between binary activity vector and binary behavior vector as described previously^10^. Statistical testing was conducted by comparing the real value to a distribution of values generated from 3000 randomly shuffled activity vectors derived from the same cell. The statistical significance level (P = 0.05) was corrected for multiple comparisons using the Bonferroni method.

The persistence of social cells across sessions was analyzed as described previously^23^ (**Figure 2G-H**). We first identified a population of cells that belonged to the functional category of interest (i.e., SCs or nonSCs) in the reference session N and then tracked the functional category of each cell in the subsequent session N+1. The likelihood of becoming a cell in each functional category in session N+1 is expressed as the probability divided by the number of cells considered in session N.

## Statistics

Data are expressed as mean ± SEM unless otherwise stated. Statistical tests were conducted using GraphPad Prism version 9 (GraphPad Software). Unpaired and paired 2 groups were compared by two-sided Mann–Whitney test and Wilcoxon matched-pairs signed rank test, respectively. More than 2 groups were compared by analysis of variance (ANOVA) with post-hoc Holm–Sidak tests. Statistical significance was defined as *P* < 0.05. Exact *P* values are shown unless *P* < 0.0001.

## Funding

This work was supported by KAKENHI from JSPS, 19H04942, 20H03550 and 23H02668 to M.S., and 16H06316, 21H04813, 23H04233 and 23KK0132 to T.T.; Japan Agency for Medical Research and Development, JP21wm0425011; Japan Science and Technology Agency, JPMJMS2299, JPMJMS229B; Intramural Research Grant (30-9) for Neurological and Psychiatric Disorders of NCNP; The Takeda Science Foundation; Taiju Life Social Welfare Foundation to T.T.

## Author contributions

Conceptualization, M.S. and E.T.N.O.; methodology, M.S. and E.T.N.O; investigation, E.T.N.O.; formal analysis, M.S., E.T.N.O. and S.F.; visualization, M.S., E.T.N.O. and S.F.; writing—original draft: M.S. and E.T.N.O.; writing—review & editing: M.S.,

E.T.N.O. and T.T.; funding acquisition, M.S. and T.T.; supervision, M.S. and T.T.

## Competing interest

The authors have no conflict of interest.

---

**Figure supplement 1.**
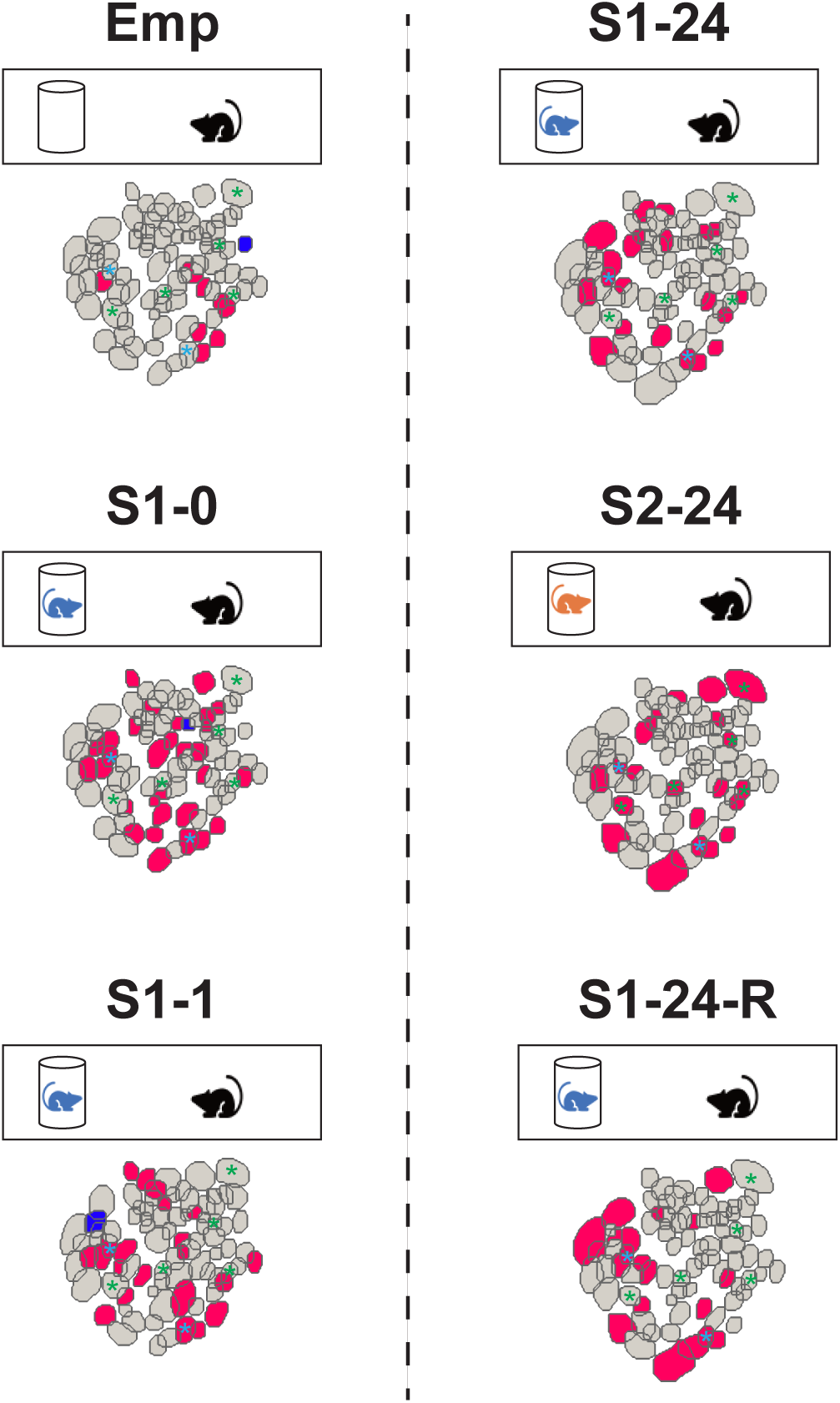
Example maps of Social-ON and Social-OFF cells imaged during each session of the SRM task. Social-ON cells and Social-OFF cells are shown in red and blue, respectively. Repeated interactions with S1 activated different combinations of cells during each session. Green asterisks indicate example cells that were Social-ON cells only during the S2–24 session. Blue asterisks indicate Social-ON cells throughout all social interaction sessions.

**Figure supplement 2.**
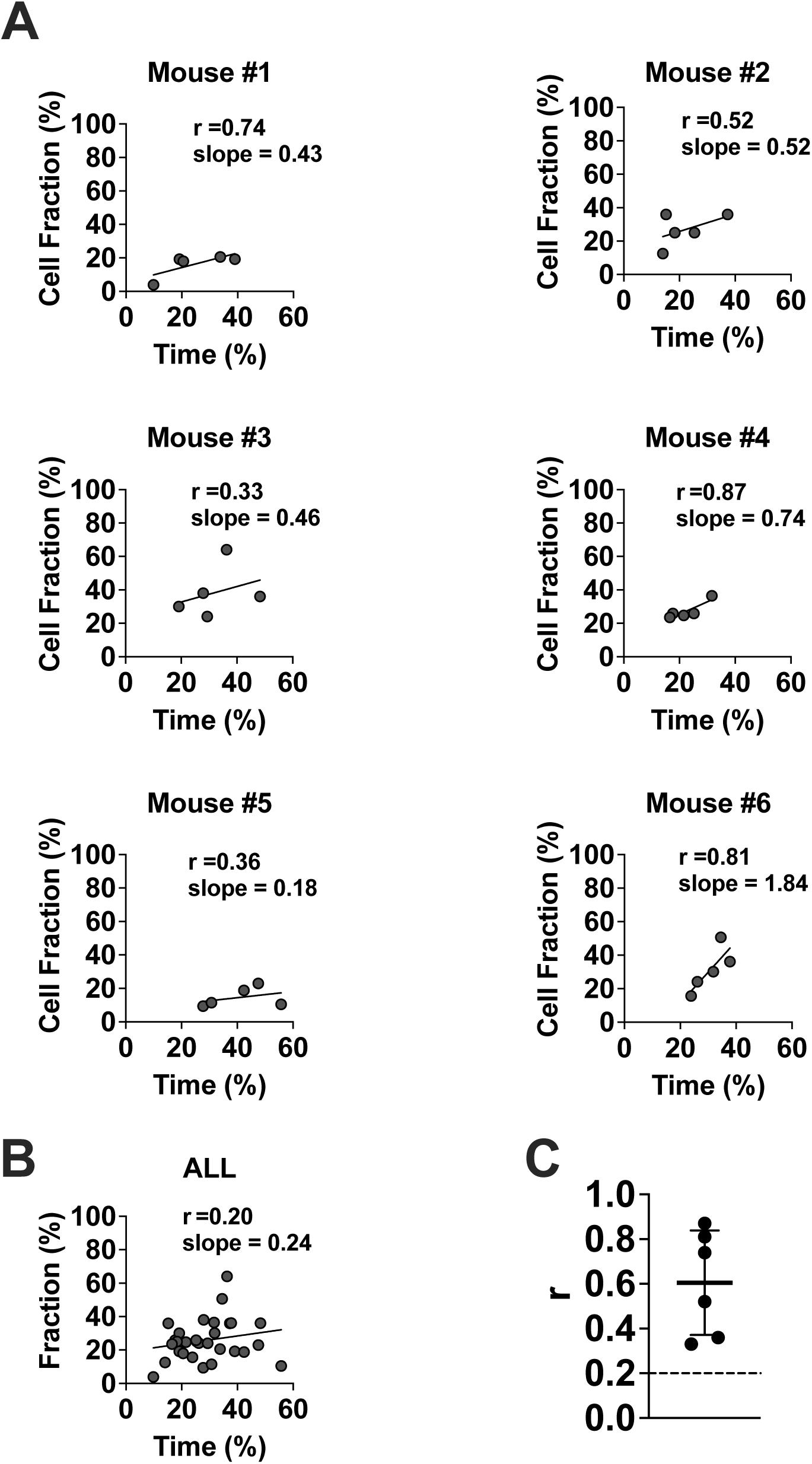
Relationship between the fraction of SCs and the total length of social interaction for each subject mouse. A. Correlations between the fractions of SCs and the total time spent investigating the social target for each subject mouse (n = 5 sessions each). **B.** Scatter plot showing the relationship between the fractions of SCs and the time spent investigating the social target for all six mice (n = 30 sessions). **C**. Linear regression coefficients for individual mice (mean ± SD of n = 6 mice). The dashed line indicates the correlation coefficient calculated from summary data for all mice.

**Figure supplement 3.**
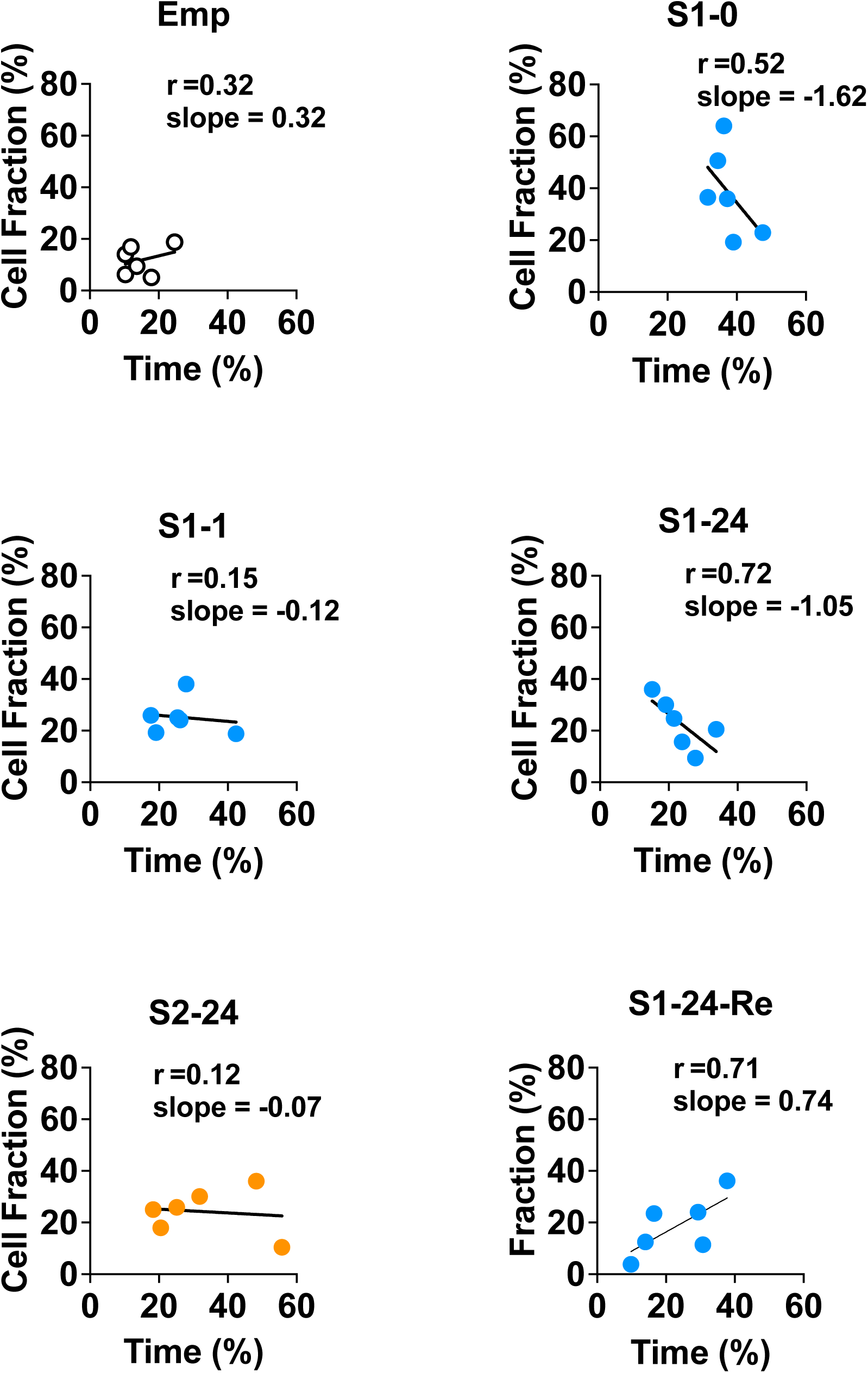
Relationship between the fractions of SCs and the total length of social interaction during each session. n = 6 mice each.

**Figure supplement 4.**
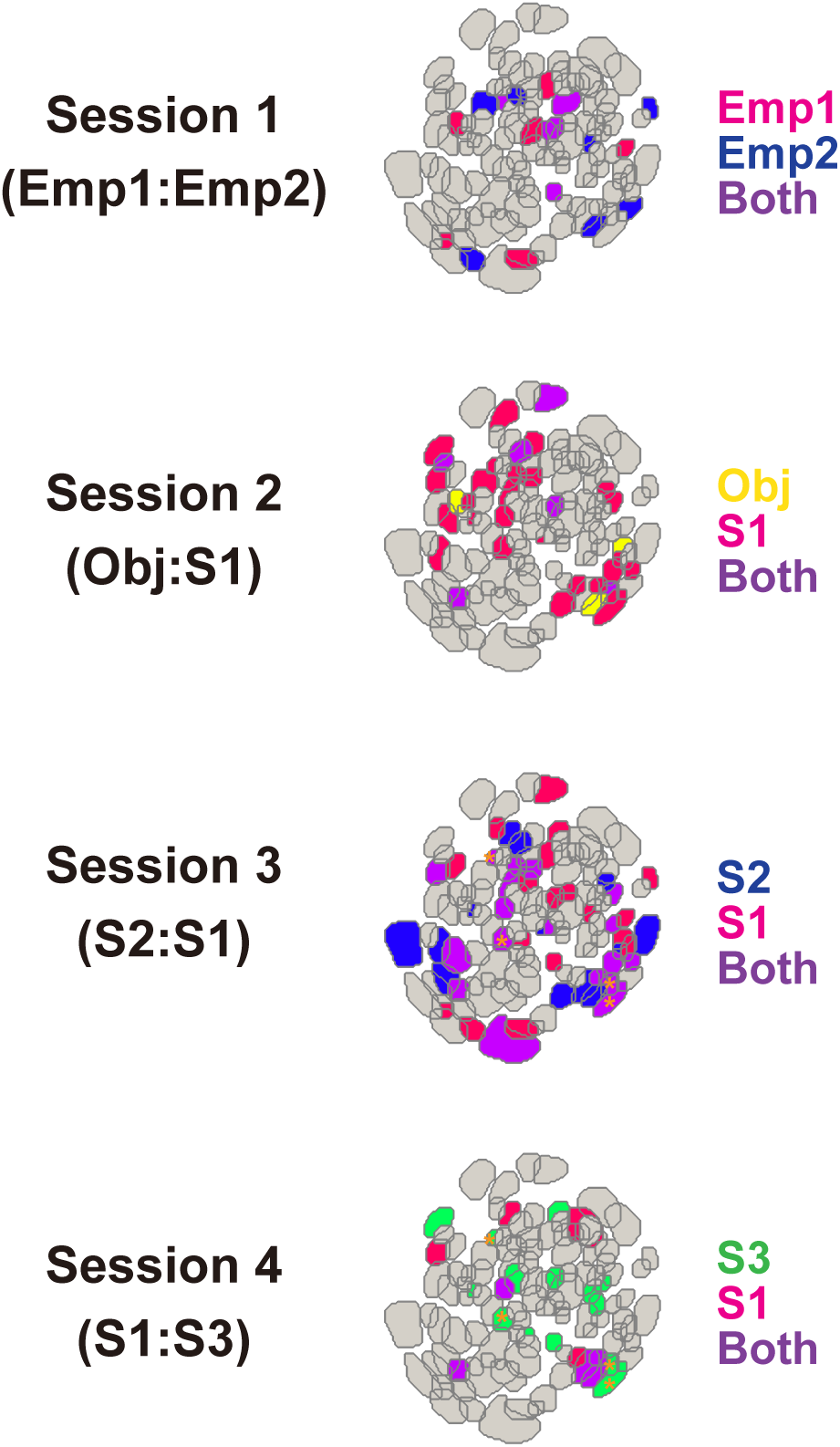
Example maps of SCs imaged during each session of the LCSD task. Colors indicate cells that responded to the targets shown on the right-hand side in corresponding colors. Cells with orange asterisks in the maps for Sessions 3 and 4 are examples that responded to both S1 and S2 in Session 3 and to novel S3 in Session 4.

## Notes

### Competing Interest Statement

The authors have declared no competing interest.

